# Identifying Impacts of Contact Tracing on Epidemiological Inference from Phylogenetic Data

**DOI:** 10.1101/2023.11.30.567148

**Authors:** Michael D. Kupperman, Ruian Ke, Thomas Leitner

## Abstract

Robust sampling methods are foundational to inferences using phylogenies. Yet the impact of using contact tracing, a type of non-uniform sampling used in public health applications such as infectious disease outbreak investigations, has not been investigated in the molecular epidemiology field. To understand how contact tracing influences a recovered phylogeny, we developed a new simulation tool called SEEPS (Sequence Evolution and Epidemiological Process Simulator) that allows for the simulation of contact tracing and the resulting transmission tree, pathogen phylogeny, and corresponding virus genetic sequences. Importantly, SEEPS takes within-host evolution into account when generating pathogen phylogenies and sequences from transmission histories. Using SEEPS, we demonstrate that contact tracing can significantly impact the structure of the resulting tree, as described by popular tree statistics. Contact tracing generates phylogenies that are less balanced than the underlying transmission process, less representative of the larger epidemiological process, and affects the internal/external branch length ratios that characterize specific epidemiological scenarios. We also examined real data from a 2007-2008 Swedish HIV-1 outbreak and the broader 1998-2010 European HIV-1 epidemic to highlight the differences in contact tracing and expected phylogenies. Aided by SEEPS, we show that the data collection of the Swedish outbreak was strongly influenced by contact tracing even after downsampling, while the broader European Union epidemic showed little evidence of universal contact tracing, agreeing with the known epidemiological information about sampling and spread. Overall, our results highlight the importance of including possible non-uniform sampling schemes when examining phylogenetic trees. For that, SEEPS serves as a useful tool to evaluate such impacts, thereby facilitating better phylogenetic inferences of the characteristics of a disease outbreak. SEEPS is available at github.com/MolEvolEpid/SEEPS.

## 1. Introduction

The growth and prevalence of communicable diseases, in which a human individual transmits a pathogen to another individual, without an intermediate vector or reservoir, has led to the development of a variety of detection and surveillance strategies. With the notable exception of zoonotic spillover events, each infection can be attributed to another, older, infection. This basic insight led to the remarkable development of contact tracing as a core method to efficiently identify closely related infections (Centers for Disease Control, 1986; Giesecke et al., 1991; Hethcote and Yorke, 1984), resulting in significant contributions to public health (Ramstedt *et al*., 1990). While contact tracing is an efficient method to prevent future infections by tracing the contacts of an index case, who may already be infected or not, and informing them about preventative measures to hinder further spread, it is also an effective method to collect samples from infected close contacts to an index case. Indeed, many pathogen sequences from outbreaks and larger epidemics have been collected via contact tracing. For example, in the recent SARS-CoV-2 epidemic, contact tracing was extensively used to collect data (Turcinovic *et al*., 2022). Generally, contact tracing has also been evaluated in mathematical models (Höhna *et al*., 2011; Müller and Kretzschmar, 2021), but there remains little knowledge on how contact tracing specifically may impact and interact with genetic sequence data analyses.

A standard form of contact tracing is “iterative contact tracing”, in which an initial index case is interviewed to identify contacts which may be infected. Identified contacts are tested, and the interview process is repeated for contacts with positive test status. The non-random nature of samples collected via contact tracing raises fundamental questions about the nature of the data that typically is used for phylogenetic reconstruction of pathogen epidemics; how robust are mathematical assumptions made about the collection of data in practice, and how significant are deviations from these assumptions in real data? While the contact network that pathogens spread across can be informative of the pathogen’s phylogeny (Giardina *et al*., 2017), it remains largely unknown how sampling with contact tracing impacts the observable phylogeny.

An alternative perspective on data collected from contact tracing is that samples obtained are the result of biased sampling (Frost *et al*., 2015). Similar issues of statistically biased sampling have been partially addressed in phylogeographic studies, where bias comes from over/under-representation of sampling in some regions, municipalities, or communities (Liu *et al*., 2022). Attempts to address this usually follow one of two strategies: perform a subsampling of the available database to find a maximally diverse subset (Townsend and Lopez-Giraldez, 2010; Townsend and Leuenberger, 2011; Dornburg *et al*., 2019; Marini *et al*., 2022), or select models that are more robust against sampling biases (Heath *et al*., 2008; Hall et al., 2016; Layan et al., 2023). The practical significance of incomplete taxon sampling has been extensively investigated and discussed (Rosenberg and Kumar, 2001; Zwickl and Hillis, 2002; Rosenberg and Kumar, 2003; Hillis et al., 2003; Heath et al., 2008; Mavian et al., 2020). While these investigations are conceptually helpful and insightful, little work has been done to understand how incomplete sampling due to contact tracing impacts the observed phylogeny in detailed simulations.

Several simulators have been proposed for generating detailed pathogen phylogenies, such as FAVITES (Moshiri *et al*., 2019), BEAST2 (Bouckaert *et al*., 2014), and PopART IBM (Pickles *et al*., 2021). However, none include contact tracing or similar sampling methods. FAVITES comes close in offering a sampling method weighted by the number of transmission events, which results in sampling towards more interconnected individuals rather than following transmission links. This method cannot completely mimic the feedback that contact tracing can have on the simulation by optionally removing sampled taxa, which can be captured within SEEPS.

A motivating real-world example to consider is the spread of HIV-1 circulating recombinant form 1 (CRF01) in Europe (Figure 1). HIV-1 CRF01 was originally introduced in Southeast Asia from Africa (Gao *et al*., 1996; McCutchan et al., 1996), and later spread from there to other parts of the world, including Europe (Hemelaar *et al*., 2020). Thus, the available HIV-1 CRF01 sequences from Europe cannot be strongly influenced by contact tracing as they are not closely related within Europe, nor were there cross-border coordinated sampling efforts. In contrast, a Swedish HIV-1 CRF01 outbreak among injection drug users in 2007-2008 (Skar *et al*., 2011) elicited a strong public health response resulting in identifying further persons who had been in contact with those infected with this HIV-1 variant, consequently sampling many closely related sequences. Hence, part of the resulting European HIV-1 CRF01 phylogeny comes from strong contact tracing, while the larger part does not. The two parts of the European HIV-1 CRF01 tree highlights the strong impact contact tracing may have on the tree structure, affecting both topological and branch length statistics.

**Fig. 1:**
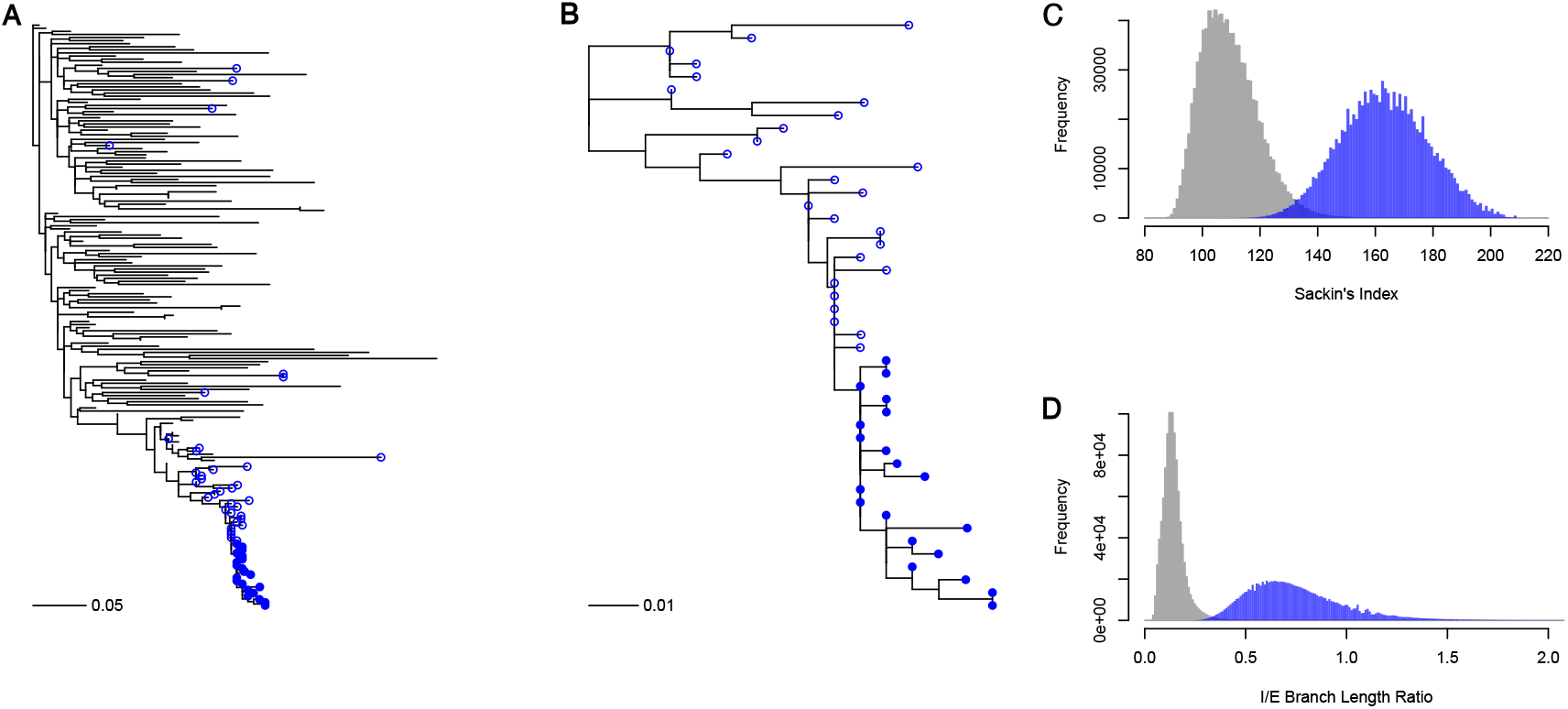
Contact tracing induces variations in phylogenetic tree structure. Panel A shows the full reconstructed HIV-1 CRF01 phylogeny of sequences collected in Europe, with tips from Sweden in blue. Filled symbols denote 20 closely related samples identified through contact tracing from a known injection drug user outbreak, while unfilled symbols denote additional Swedish samples. Panel B zooms in on the bottom subtree consisting entirely of Swedish sequences. Note that the shape of this subtree is drastically different from the full tree in panel A. To quantify the difference we downsampled 20 tips randomly (without replacement) from the tree without the Swedish subtree taxa in panel A and the Swedish taxa in panel B 1,000,000 times each, and recorded both the Sackin’s index and internal to external (I/E) branch length ratios in panels C and D, respectively. The blue distributions are from the Swedish subtree in panel B and the gray distributions from the full European tree without the Swedish subtree taxa. Comparing the distributions with a Kolmogorov-Smirnov test showed very different distributions: D = 0.964, p*<* 10^−16^ for Sackin’s index and D = 0.988, p*<* 10^−16^ for the I/E branch length ratios. Trees were inferred by maximum-likelihood under a GTR+I+G substitution model (Guindon *et al*., 2010). Scale bars in panels A and B are in units of substitutions/site.

To directly address questions about the significance of contact tracing, we developed a new simulation suite in R called SEEPS (Sequence Evolution and Epidemiological Process Simulator).

## 2. Methods and data

### 2.1. SEEPS overview and availability

Building off the agent-based HIV-1 model in Kupperman *et al*. (2022), we developed SEEPS, an end-to-end modern and modular simulator for investigating the connection between evolutionary and epidemiological mechanisms. Written in R (R Core Team, 2022), SEEPS is a flexible and extensible framework for simulating phylodynamic and evolutionary processes at a population level (with the capability to simulate the within-host evolutionary processes at the same time). A general schematic of SEEPS is shown in Fig 2. SEEPS uses a stochastic forward simulation that tracks the transmission history of the entire simulation (including non-sampled individuals) and maintains a list of active individuals that are capable of generating new offspring. SEEPS stores the entire transmission history, allowing for contact tracing to be applied on top of the transmission history, just as in reality.

**Fig. 2:**
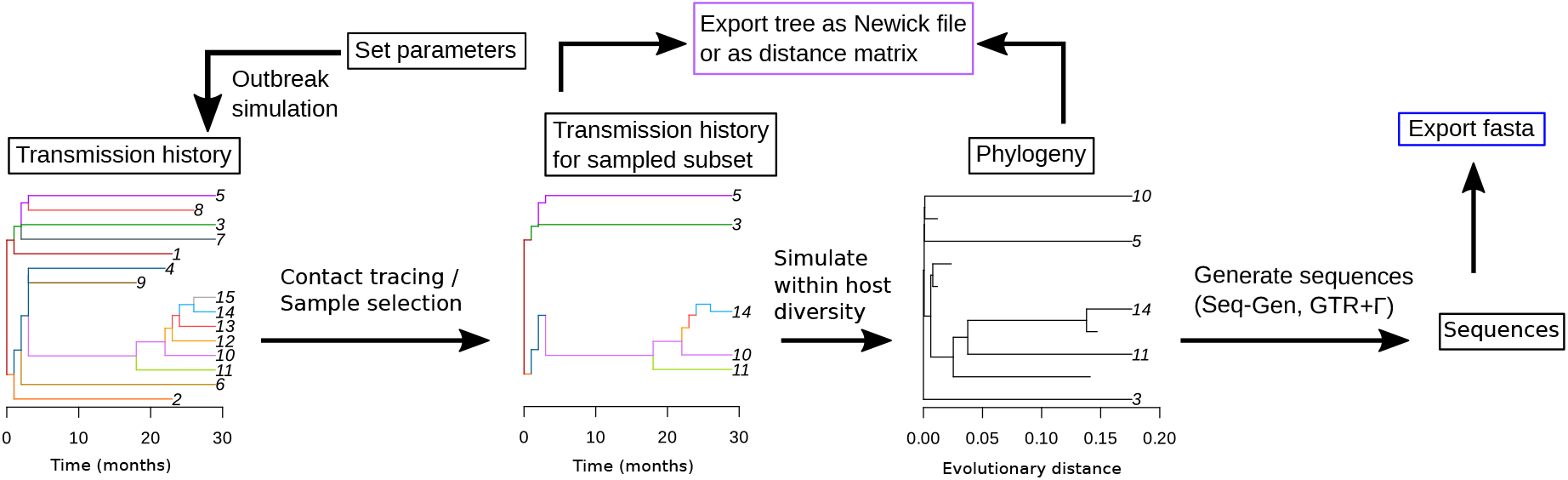
General workflow for the SEEPS package. Arrows denote functionality provided by SEEPS. Blue and purple boxes denote exportable data. Black boxes denote internal states or data available for manipulation.

All simulations in the present study were performed in SEEPS v0.2.0 available at github.com/molEvolEpid/SEEPS. Code associated with specific analyses is available at github.com/molEvolEpid/ContactTracingForPhylogenies. Below, we provide a brief guide for using SEEPS.

### 2.2. Specific steps in SEEPS

#### 2.2.1. Simulating transmission dynamics, including within-host evolution

An experiment in SEEPS begins by simulating transmission events in a population with a user-defined offspring distribution and generation time. Individuals in SEEPS are considered *active* if they are capable of generating secondary infections. The transmission history is recorded. Infected individuals are sampled at user-defined time points before being removed from the simulation, forming the sampled transmission tree.

SEEPS offers a module for simulating the within-host viral evolution for each infected individual using a coalescent process from Lundgren *et al*. (2022).

#### 2.2.2. Outputs from SEEPS

SEEPS exports trees in Newick format for use in other standard phylogenetic analysis software. Sequence simulation is available with a GTR+I+Γ model using Seq-Gen (Rambaut and Grass, 1997) and the PhyClust R package (Chen, 2011). Distance matrix representations of the data are also available for export, either using cophenetic distances for trees, or pairwise evolutionary distances (such as TN93) for sequences.

#### 2.2.3. Simulating contact tracing

SEEPS captures the fundamental aspect of iterative contact tracing, where each positive contact is discovered with probability 0 ≤ *p* ≤ 1. If *p* = 0, there is no contact tracing, while *p* = 1 corresponds to perfect contact tracing. It is similar to the popular breadth-first-search algorithm (Lee, 1961), but with the variation that the discovery of edges is randomized with a prescribed failure probability 1 − *p*.

Algorithmically, an initial index case is randomly discovered in the population, and all direct contacts (secondary infections and the source itself) are identified in the transmission history. Then, each contact is *independently* discovered at probability *p* ∈ [0, 1]. The discovered individuals are added to a list of discovered individuals, and the identify-and-probabilistically-discover process is repeated for each newly discovered individual. Importantly, in agreement with the fact that real-world contact tracing may prevent future infections, we assume that persons identified through contact tracing do not transmit further from the time of diagnosis due to immediately being put on effective antiviral treatment.

### 2.3. Settings of SEEPS simulations used in the study

We simulated transmission dynamics assuming that the length of each infection is a uniform distribution between one and three years and the expected number of lifetime transmissions is *R*_0_. Following Graw et al. (2012), we assumed that the transmission potential was 20-fold higher in the first three months of infection to reflect the higher rate of transmission during the acute infection phase. The simulation time step was 1 month.

After prescribing the population dynamics (here, exponential growth or constant size), samples were taken at fixed time points at a specified contact tracing probability.We saved both the reduced transmission history, where unsampled tips were removed and any internal nodes of order two were collapsed, and the complete transmission history for the samples. The complete transmission history for the sampled individuals was used to obtain a phylogeny by simulating a coalescent process along the transmission history (Lundgren *et al*., 2022). Within-host diversity was modeled assuming an expected maximum number of transmitted lineages with each new infection at *α* = 5, and within-host diversification rate of number of lineages *β* = 5 per day, as in Romero-Severson et al. (2016). Individuals in the transmission history that are not sampled are simulated where needed to ensure that the entire evolutionary history of the samples is correct with respect to transmission bottlenecks and diversification level until the next transmission event. This process can introduce additional tips into the phylogeny that do not correspond to sampled individuals. These were removed in our main analyses, but are available within SEEPS for inspection (as shown in Fig S3).

### 2.4. Tree statistics

As phylogenetic trees are complex objects, there are many statistics, indexes, and measures that have been proposed for analyzing trees (Fischer *et al*., 2021). We considered Sackin’s index (Sackin, 1972; Shao and Sokal, 1990) using the R package treebalance (Fischer *et al*., 2021) to assess topological effects, and the internal/external (I/E) branch length ratio to assess branch length effects.

Sackin’s index measures the imbalance of a tree. It is maximized in a caterpillar (ladder-like) topology. In the absence of contact tracing, i.e., with uniform random sampling, we expect the Sackin’s index to be low, because then the topology would be informed by random ancestral relationships during the initial exponential growth phase. In the presence of contact tracing, we expect to primarily recover recent information about the transmission history in active epidemiological clusters. While such clusters also are linked together by ancestral relationships, if the contact tracing is good, we expect the majority of the tree to reflect recent transmission events.

We used parsimony to assess how representative a sample of taxa is of a larger epidemic. We did this by sampling twice in an epidemic and compared how representative the second sample was of the phylogeny obtained from the first sample. The taxa of the first sample were labeled “A” and the taxa of the second sample “B”. Given the phylogeny of both “A” and “B” labeled taxa, we calculated the number of “A” → “B” transitions, i.e., the parsimony score of label transitions. If uniform random sampling was performed, we expect the parsimony score to be high. If contact tracing was performed, we expect each group of taxa to contain more cluster-like relationships, which would be more informative about local spread but not the entire epidemic. Thus, because more taxa are closely related when contact tracing has occurred, fewer “A” → “B” transitions are required, and the resulting tree would have a lower parsimony score.

The I/E ratio is informative of the recent evolutionary relationship between taxa, as well as the overall tree structure (Giardina *et al*., 2017). In the absence of contact tracing, we expect the ratio to be low, as many external branches will connect the taxa back to an ancestral event in the outbreak phase. In contrast, if there is contact tracing, we expect the ratio to be high as the most recent common ancestor between two taxa in an identified cluster will be much more recent.

### 2.5. HIV-1 CRF01 European sequence data

Data from the European HIV-1 CRF01 epidemic was extracted from the LANL HIV database (hiv.lanl.gov). GenBank accession numbers for all sequences used in this study are available at github.com/molEvolEpid/ContactTracingForPhylogenies. The data consisted of 34 *env* V3 region sequences (approx. 300 nt) from an intravenous drug user (IDU) outbreak in Stockholm, Sweden, in 2006-2007 (Skar *et al*., 2011) and 155 European sequences from 2003-2007 (including 23 additional Swedish sequences not involved in the IDU outbreak and 132 sequences from 12 other countries). The entire European HIV-1 CRF01 tree was reconstructed using PhyML v3 under a GTR+I+G model by both NNI and SPR search (Guindon *et al*., 2010).

Using SEEPS, we simulated 110,000 and 3,520,000 sequences for the EU and Swedish outbreaks respectively with varying levels of contact tracing. Sampling dates were selected by sampling the distribution of sample years that the true sequences were taken from. We then computed the tree statistics for each simulated outbreak. To compare the simulated sample distributions against the bootstrap distributions computed from the real data, we used the two-sample Kolmogorov-Smirnov test and the absolute relative difference of means. As these simulations were large, we refrained from reporting a p-value, but instead reported the test statistic as evidence for how close the simulated distribution were to the real data. For both test statistics, a value closer to zero implies that the simulated distribution was closer to the real data.

## 3. Results

### 3.1. Contact tracing impacts general tree structures

Highly successful contact tracing results in strongly connected clusters being identified. In contrast, low amounts of contact tracing results in identifying small clusters that are loosely related in the past. Fig S3 shows examples of simulated trees under high and low contact tracing probabilities. These examples show both the complete transmission histories and trees derived after sampling, which would be the typical case in real-world data. Importantly, sampling by contact tracing can have profound impact on overall tree structure. Thus, assuming random sampling (which is typically done in phylodynamic inferences) may give a very different impression of what appeared to have happened.

### 3.2. Contact tracing makes trees less balanced

To evaluate the cladistic impact of contact tracing, we first measure Sackin’s index for a collection of taxa taken at a single time point, known as cross-sectional sampling. We simulated 1,000 outbreaks followed by a constant population size for 0 to 10 years (in one year increments), with *R*_0_ uniformly distributed between 1.5 and 5. For each outbreak, we sampled either 15 or 50 taxa, with contact tracing performed at either high (*p* = 0.9) or low (*p* = 0.1) levels. In total, we generated 22,000 transmission trees and 22,000 phylogenies.

We found no clear correlation between Sackin’s index and *R*_0_ and no effect of the number of years after the outbreak phase in when a sample was taken. The simulation of the transmission history resulted in an average Sackin’s index close to what could be predicted from a Yule model (Kirkpatrick and Slatkin, 1993) when contact tracing performance was low (Fig 3). Conversely, when contact tracing was high (*p* = 0.9), Sackin’s index became elevated above the Yule expectation. Adding within-host diversification increased Sackin’s index only slightly for both low and high levels of contact tracing. In all configurations, the sampled trees include a Sackin’s index close to the minimal possible value for the number of sampled taxa (15 or 50) (Fischer, 2021).

**Fig. 3:**
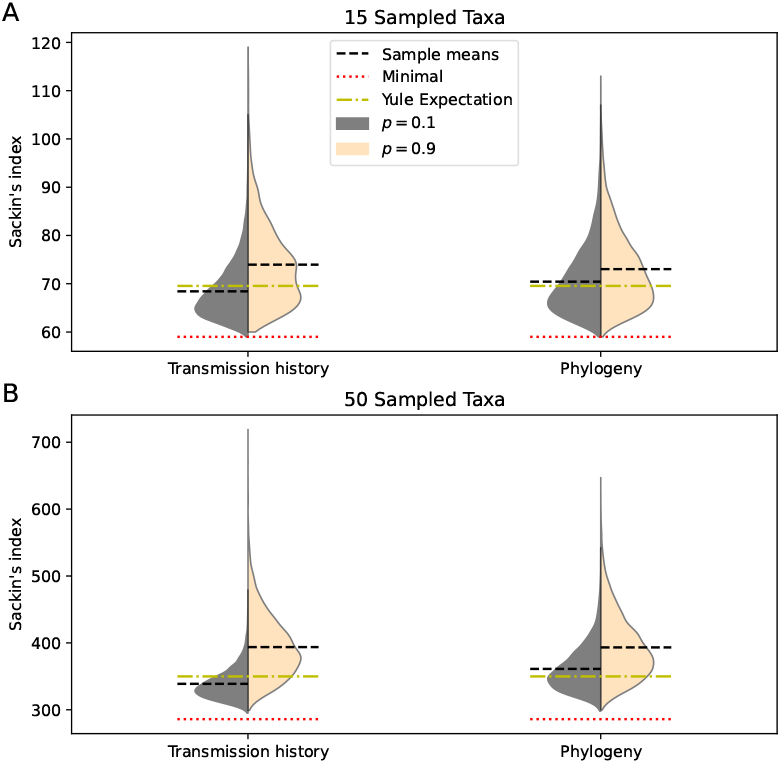
Violin plots of the distribution of Sackin’s index with low (*p* = 0.1) and high levels of contact tracing (*p* = 0.9). Horizontal reference lines are added at the minimal value of Sackin’s index (red), the expected value of Sackin’s index under a Yule model (yellow), and the sample mean (black). Adding contact tracing increases Sackin’s index, regardless of whether the transmission history tree or the sampled phylogenetic tree is considered.

### 3.3. Trees based on contact tracing in a small sample do not represent the larger epidemic

Since we now know that contact tracing biases trees to be more unbalanced, this raises concern about how representative a phylogeny would be of the greater epidemic. Thus, next we assessed how representative a second sample would be of an earlier sample from the same epidemic.

We simulated outbreaks under varying *R*_0_ to an effective population size of 1,000 infections and sampled N active infections as soon as the effective population surpassed 900 active infections. We let the population replace the removed infections while simulating forward for 3, 24, or 120 months. We then drew another N taxa, for a total of 100 sampled taxa. The growth rate parameter *R*_0_ took discrete values of 1.1, 1.5, 3, 5, or 10.

We used a parsimony score to report the number of “transitions” that were required to render the first sample labels into the second sample labels. Thus, a higher score would indicate a more similar tree, while a smaller score would indicate a more different tree.

We found a strong relationship between the mean parsimony score and both contact tracing and the length of the inter-sampling period (Fig 4 A-E). Increased contact tracing decreased the parsimony score, indicating that the two samples represented different parts of the total epidemic. *R*_0_ only weakly influenced the relationship between parsimony scores and contact tracing performance; *R*_0_ primarily impacted the parsimony score when contact tracing performance was high by increasing the variance. Setting *R*_0_ = 10 indicated that the variance increased after approximately *p* = 0.5, while the decrease in variance occurred close to *p* = 1 when *R*_0_ was close to 1. Fig S1 and S4 shows additional results to provide a more complete picture of this effect.

**Fig. 4:**
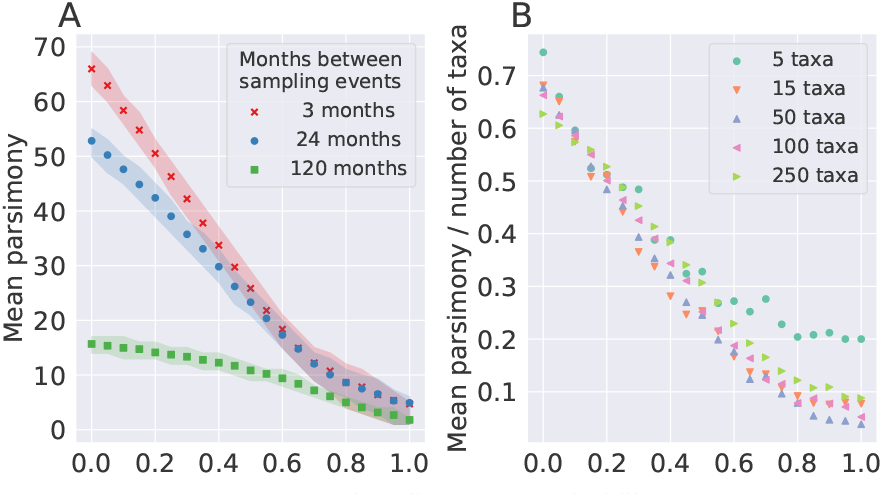
Parsimony distributions were strongly related to contact tracing level *p* and weakly to sample size. Here, *R*_0_ = 1.5. Other values of *R*_0_ are shown in Fig S4. In panel A, a sample of 100 taxa was obtained at the end of the exponential growth phase, and compared against another sample of 100 taken 3, 24, or 120 months later. The shaded regions denote the symmetric inner 50% of the results. In B, the experiment is repeated for the 3-month interval for *N* = 5 to *N* = 250 taxa. The parsimony score was normalized against the number of taxa in the sampled tree.

The sample size (number of taxa) barely influenced the parsimony score (Fig 4B). For a “small sample” of five taxa, the lowering effect of contact tracing on the parsimony score became diminished because the small tree size limits its range.

### 3.4. Mean internal to external branch length ratio is affected by contact tracing

The mean internal to external (I/E) branch length ratio quantifies the recent evolutionary relationship between sampled taxa. If the ratio is small, this suggests that the taxa are not recently related. If the ratio is large, then the samples are recently more related, suggesting the possibility of an epidemiologically significant cluster. Previous work (Giardina *et al*., 2017) suggests that the branch length ratio can be informative about possible recent outbreaks in a population, but the impact of contact tracing on the I/E ratio has not been evaluated.

Using the same simulated data we examined for Sackin’s index, we computed the I/E branch length ratio for each phylogenetic tree (Figure 5). While the *R*_0_ growth rate of the outbreak had some impact on the mean I/E branch length ratio, the presence or absence of contact tracing amplified the effect of when sampling occurred relative to the epidemic outbreak on the I/E branch length ratio. *R*_0_ was influential only when it was low; epidemics at *R*_0_ *>* 2.5 taken at the same time point had similar I/E branch length ratios, whereas the I/E branch length ratios increased when *R*_0_ was *R*_0_ *<* 2.5. Interestingly, the I/E branch length ratio was less sensitive to contact tracing immediately after the peak of the outbreak. At low contact tracing (*p* = 0.1), the I/E ratio was never able to rebound past the initial outbreak signal. In contrast, at high contact tracing (*p* = 0.9), the samples taken three years after the end of the exponential growth phase had similar I/E branch length ratios to the samples taken immediately after the peak of the outbreak. Thereafter, the I/E branch length ratio statistic continued to grow with time.

**Fig. 5:**
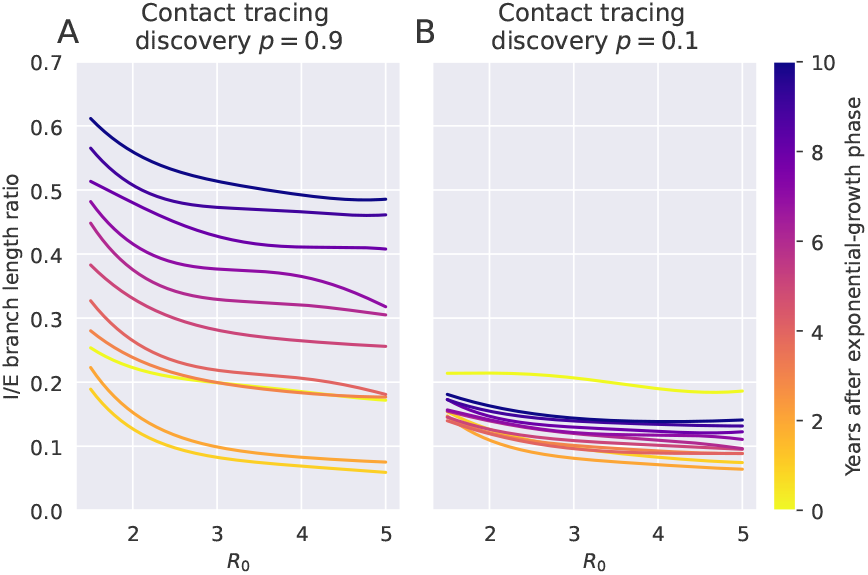
Estimated mean of the internal to external (I/E) branch length ratios as a function of *R*_0_, stratified by number of years after the peak. The mean trend line was estimated using a Gaussian process model (radial basis function kernel, *α* = 3 *×* 10^−5^, *l* = 3) (Pedregosa *et al*., 2011). The color of the trend line indicates when sampling was performed after the epidemic exponential growth had ended. The I/E ratio interpretation depends on both *R*_0_ and contact tracing level.

This suggests that some amount of contact tracing early in an epidemic, enough to find a recent nearby infection, is necessary to recover a time signal from the internal branches and indicate the age of the outbreak.

### 3.5. Contact tracing can be observed in real data

Our simulations showed that contact tracing has strong effects on phylogenetic tree reconstructions, and therefore on any epidemiological inference that would be based on such trees. To tests whether we could recover the epidemiological data of our motivating example in the introduction, including the levels of contact tracing in the European and Swedish partitions, we attempted to use SEEPS to simulate the European HIV-1 CRF01 epidemic.

To estimate the level of contact tracing, we simulated epidemics in SEEPS similar to the European and Swedish outbreaks with varying levels of contact tracing and compared the I/E branch length ratios from our simulated trees to that of the real data (Figure 1). The parameter values used to generate the simulated data are shown in table S1. While we used different parameters for each outbreak, we used a similar two-phase simulation for both the European and Swedish partitions: In the first phase, we started the simulation with a single infected individual and allowed the population to grow to a small, fixed size. Once the population reached the fixed size, we let it continue at that size until the end of the phase. In the second phase, we increased the effective population size to our target value, and allowed *R*_0_ to change. To mimic the import to Europe of genetically distant lineages from Thailand, we shifted the sampling time of the European sequences forward by 18 years to reflect the lack of available sequences along the long branches that constitute the introductions into Europe. Finally, we sampled 20 individuals with varying levels of contact tracing discovery probability according to the sample years of the EU and Swedish outbreaks, respectively. We then calculated the I/E branch length ratio for each simulated tree and compared the distribution against the real data distribution.

To compare simulated and real I/E ratio distributions, we computed the absolute relative difference of means 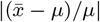 and the Kolmogorov-Smirnov test statistic *D* = sup_*x*∈[0,∞]_ |*F* (*x*) − *G*(*x*)| where *F* and *G* are the cumulative distribution functions for the simulated and the real data, respectively. For these two fitting statistics, we effectively randomized both the population size and *R*_0_ parameters, resulting in 10,000 samples to approximate the European epidemic and 320,000 samples to approximate the Swedish outbreak at each level of contact tracing probability *p*.

Both the KS statistic and the absolute relative difference of means suggested that the European data was generated in a situation with negligible contact tracing, with an upper bound on the contact tracing discovery probability of at most 10% (Figure 6). In contrast, the subsampled Swedish data were indicative of a contact tracing discovery probability of approximately *p* = 0.6. We expected that the level of contact tracing would have been very high in this intensely followed outbreak (Skar *et al*., 2011). However, two effects may have lowered the estimated contact tracing level: 1) Not all infections in the outbreak may have been sampled, which would affect the I/E ratios, and 2) subsampling the Swedish outbreak phylogeny removed over half of the taxa in each draw, which further may have lowered the estimated contact tracing level.

**Fig. 6:**
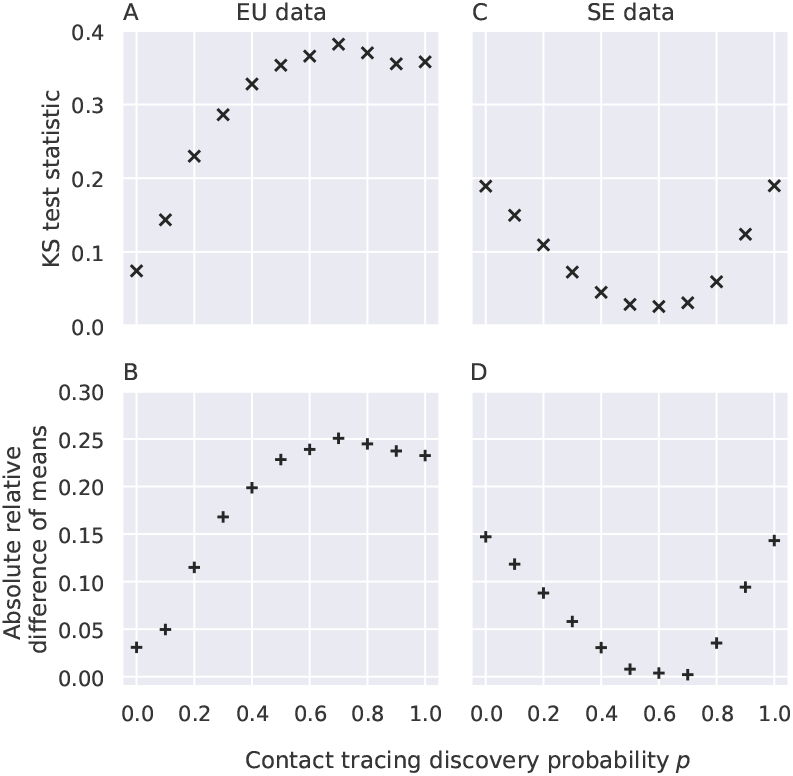
Comparison of simulated I/E branch length ratio with subsamples taken from the European epidemic and the Swedish outbreak. The contact tracing discovery probability *p* was varied from 0 to 1. Panels A and B compare simulated data against the European Union subsamples, while panels C and D compare against the Swedish subsamples.

## 4. Discussion

We developed an epidemiological and evolutionary simulator that includes contact tracing, available as a R package called SEEPS. We showed that when pathogen sequences were collected by contact tracing there can be serious impact on the resulting phylogenetic tree structure. Overall, contact tracing resulted in a phylogeny that 1) was more unbalanced, 2) was less representative of the larger epidemic, and 3) had its I/E branch length ratio differently impacted depending on when samples were taken relative to an outbreak. We then analyzed a real data set describing a known outbreak of HIV-1 CRF01 in Sweden, derived from Europe, which in turn was derived from several lineages from Thailand. This showed that SEEPS was able to simulate a fairly complicated epidemiological scenario and, importantly, also correctly detected contact tracing as it was used in Sweden.

Because sequence- and phylogeny-based approaches have the potential to reveal otherwise difficult to measure details about how pathogens spread, previous work has evaluated several tree statistics related to the branching structure and the branch lengths, such as Sackin’s index and internal to external (I/E) branch length ratios. Here, we showed that both of these classes of tree measurements are affected by contact tracing. Thus, assuming that sequences have been randomly collected, when in fact they were collected by contact tracing may severely mislead analyses and conclusions from sequence- and phylogeny-based epidemiological inferences.

While contact tracing may lead to a phylogeny that does not represent the larger epidemic - from a public health perspective, detecting superspreaders is important because they contribute more to overall disease spread. Contact tracing will be much more likely to find superspreaders than random sampling simply because they are more likely to be traced from any one of the people they infected.

SEEPS includes within-host evolution that simulates diversification under a neutral coalescent process. The within-host pathogen diversification is important to account for because it affects the observable phylogeny, which, at least for HIV, always is different from the non-observable transmission history (Graw *et al*., 2012; Romero-Severson *et al*., 2014; Giardina *et al*., 2017). However, SEEPS does not simulate the selection of escape mutants driven by the host immune system. Because selection also can cause an imbalance in the tree structure, our simulations may be on the conservative side of the impact contact tracing has on the global tree structure of an epidemic. Furthermore, if superspreaders were active, the simulations under our neutral model may show less impact of superspreading than in real epidemics.

Methods that depend on analyzing distances such as HIV-TRACE (Kosakovsky Pond *et al*., 2018) or machine learning based methods such as convolutional neural network (CNN) models (Kupperman *et al*., 2022) are inherently sensitive to the distribution of pairwise distances. Contact tracing results in samples that can be significantly closer than random. If clusters are interpreted as outbreaks, then clusters discovered by non-uniform sampling may be correctly labeled as transmission clusters, but erroneously inferred as signs of a larger outbreak. Thus, the performance of HIV-TRACE and the CNN model in detecting outbreaks may be sensitive to how samples were collected. Hence, popular analytical methods and computational tools used to trace and reconstruct epidemics need to ensure that the impact of contact tracing is not being overlooked or misinterpreted.

## 5. Acknowledgement

This work was supported by the National Institutes of Health (NIH) grants R01AI087520 and R01AI135946 to TL.

## 6. Data Availability

SEEPS is available at github.com/MolEvolEpid/SEEPS. Additional data used in this study is available at github.com/MolEvolEpid/ContactTracingForPhylogenies.

