## Supplemental materials for "Identifying Impacts of Contact Tracing on Epidemiological Inference from Phylogenetic Data"

Supplementary material

Michael D. Kupperman[1,2], Ruian Ke [1], and Thomas Leitner[*,1]

June 19, 2024

Table S1: Parameter values used to generate simulated data in SEEPS to approximate EU and Swedish outbreak samples subtree exterior/interior branch length ratio distributions. Transmission rate ratio is the proportion of the transmission rate in the first 3 months of infection, relative to the remaining average 21 months.

| Parameter | EU simulation value | SE simulation value |
|---|---|---|
| Number of trials per parameterization | 200 | 10,000 |
| transmission rate ratio | 20:1 | 1:1 (uniform) |
| Phase 1 length [Years] | 2 | 1 |
| Phase 1 $R_0$ | 3, 5 | 3, 5 |
| Phase 1 maximum effective population size | 7 | 100 |
| Phase 2 length [Years] | 2 | 5.5 |
| Phase 2 $R_0$ | 2, 3, 4, 5, 6 | 2, 3, 4, 5 |
| Phase 2 effective population size | 600, 700, 800, 900, 1000 | 25, 30, 40, 50 |
| Sampling time shift [Years] | 18 | 0 |

[1] Theoretical Biology and Biophysics group, Los Alamos National Laboratory, Los Alamos, NM, USA
[2] Department of Applied Mathematics, University of Washington, Seattle, WA, USA

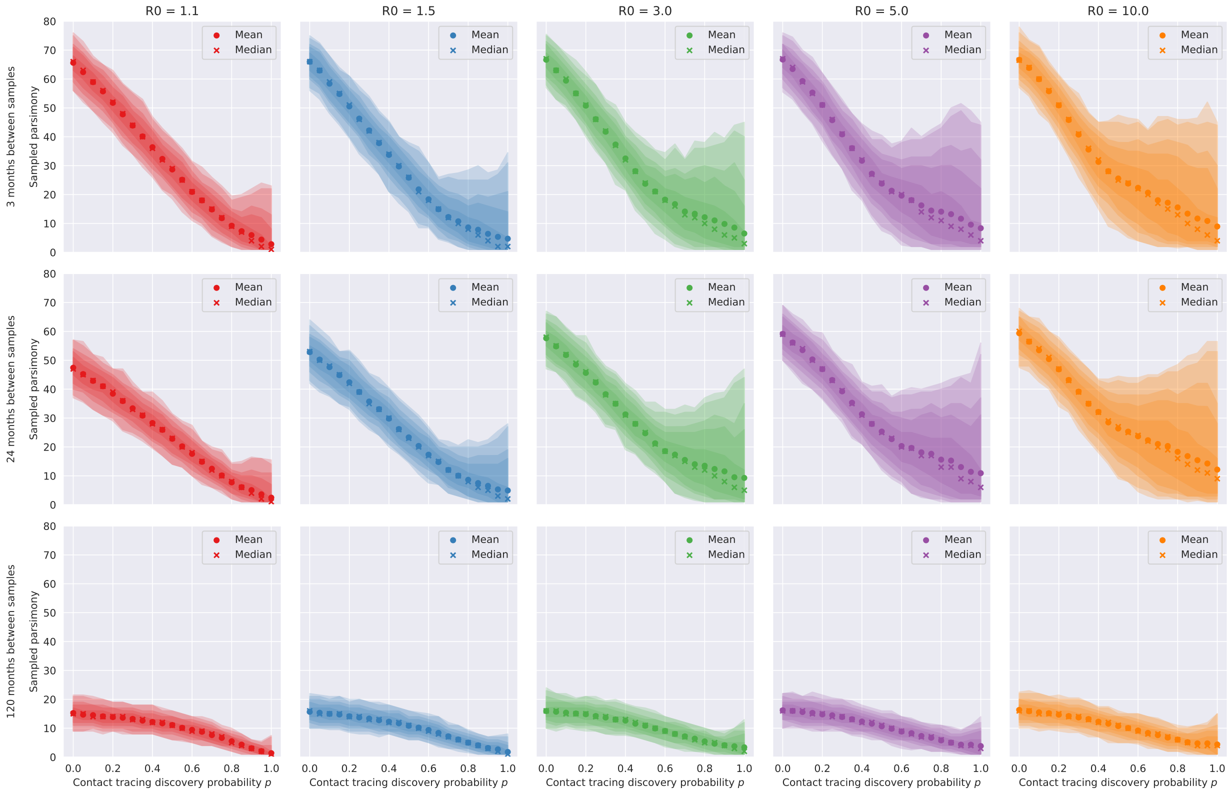


Fig. S1: More detailed representation of the parsimony distributions summarized in Fig 5. Each panel shows the sampled parsimony distributions with contours drawn at the inner symmetric 99%, 98%, 90%, 80%, and 50% percentiles respectively. We can generally identify two regimes, partitioned into a *low* amounts of contact tracing and *high* amount of contact tracing. The amount of contact tracing in our model that separates the two regimes varies as $R_0$ changes. While there is an appreciable shift in the mean parsimony score as the contact tracing parameter increases, the two regimes are primarily identified through the change in spread.

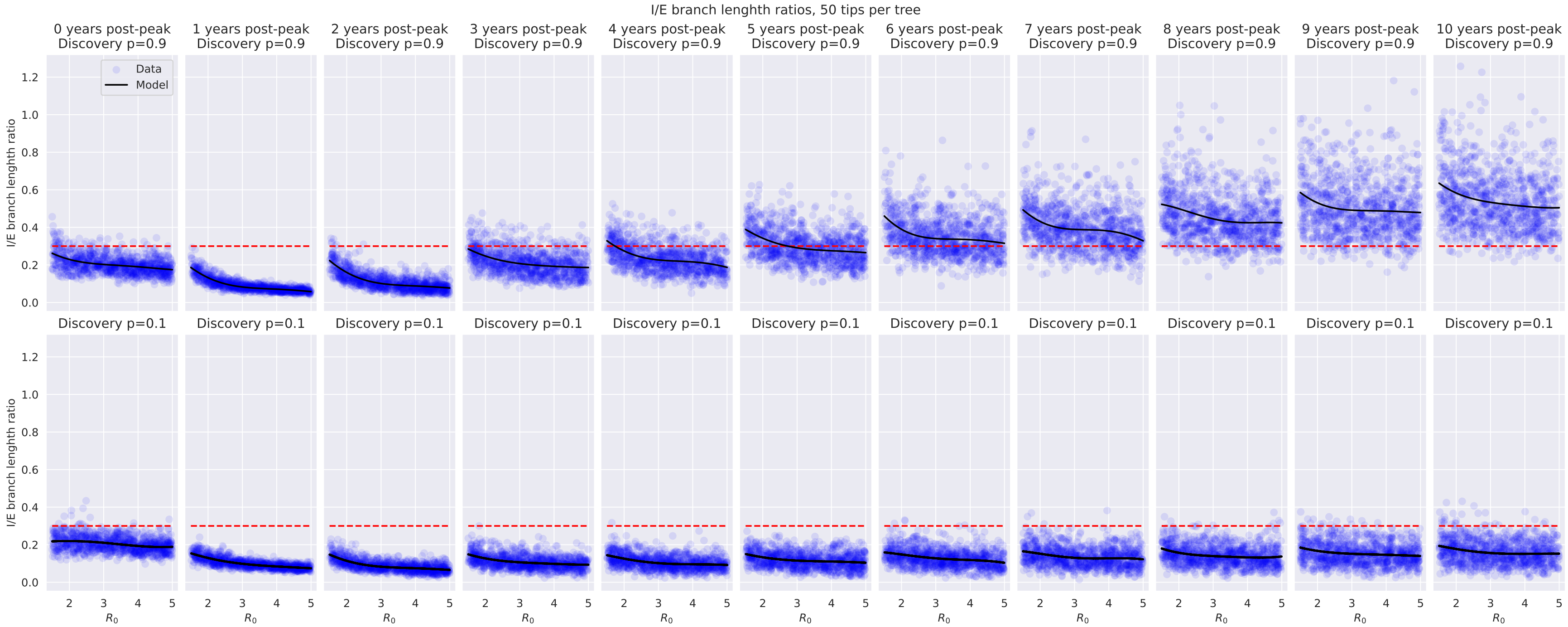


Fig. S2: Distribution of sampled I/E branch length ratios from independent samples taken between 0 and 10 years after the end of an exponentially growing outbreak, during a constant phase with $R_0$ randomized. The top row is generated with very high levels of contact tracing (discovery probability $p = 0.9$) and the bottom row is generated with very low levels of contact tracing (discovery probability $p = 0.1$). Iterative contact tracing with restarts is used in both cases. A simple regression model with a Gaussian kernel is used ($\alpha = 3 \times 10^{-5}$ with $\ell = 3$) to generate a mean trend line, which is plotted. A reference line at 0.3 is plotted. This ad-hoc threshold gives a good separation between the two resulting distributions at the 10-year mark.

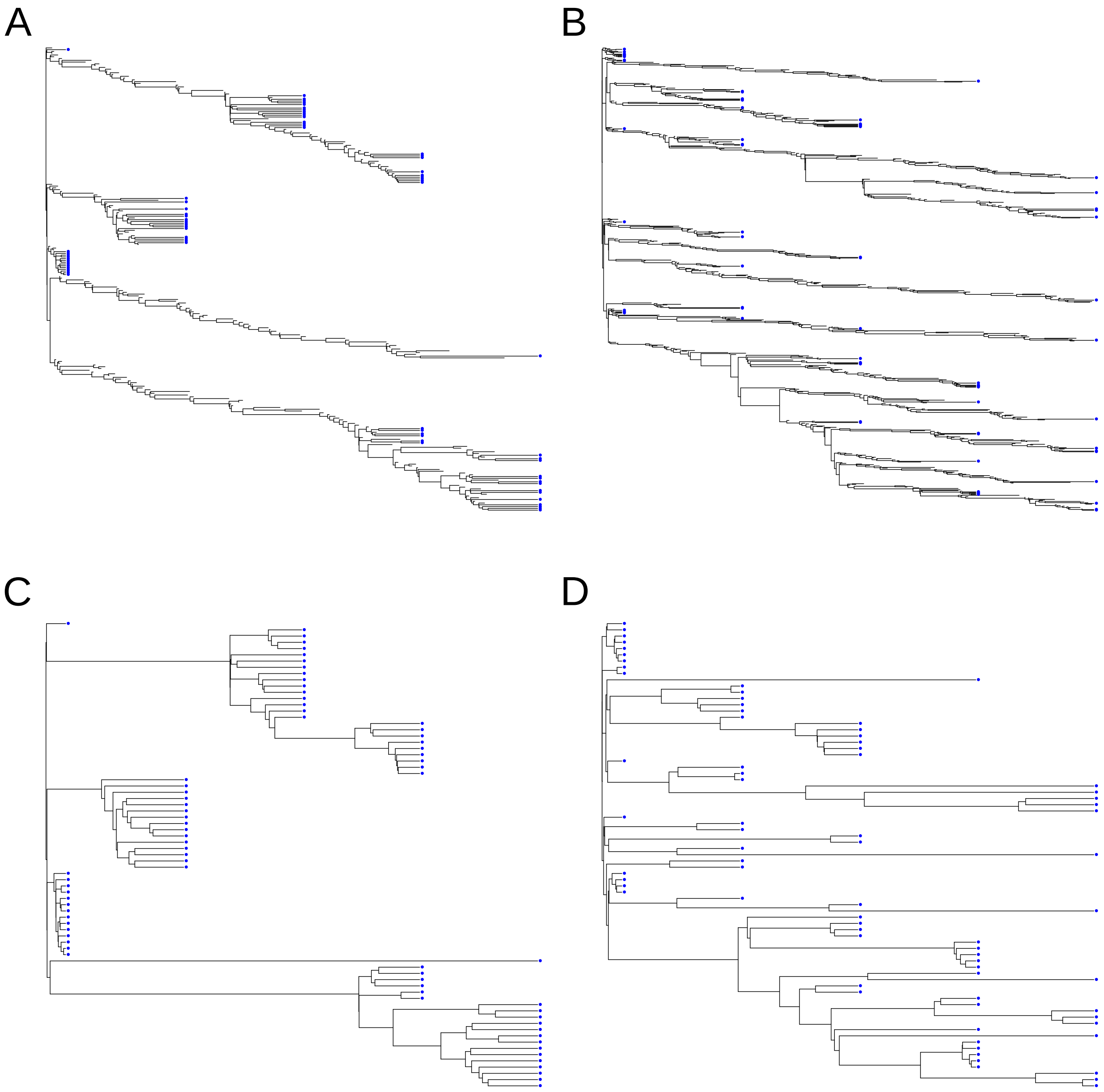


Fig. S3: Examples of how repeated sampling with different contact tracing levels can lead to visually distinct tree structures. Panel A (high contact tracing) and panel B (low contact tracing) show examples of two simulations provided by SEEPS. Note that SEEPS simulates within host diversity for all ancestors, resulting in explicit modeling of the transmission bottlenecks and the realistic population diversity. Panels C and D show only the "observable" history that could be inferred by reconstructing the ancestral relationships between the observed sequences. Sampled taxa are denoted by a blue circle.

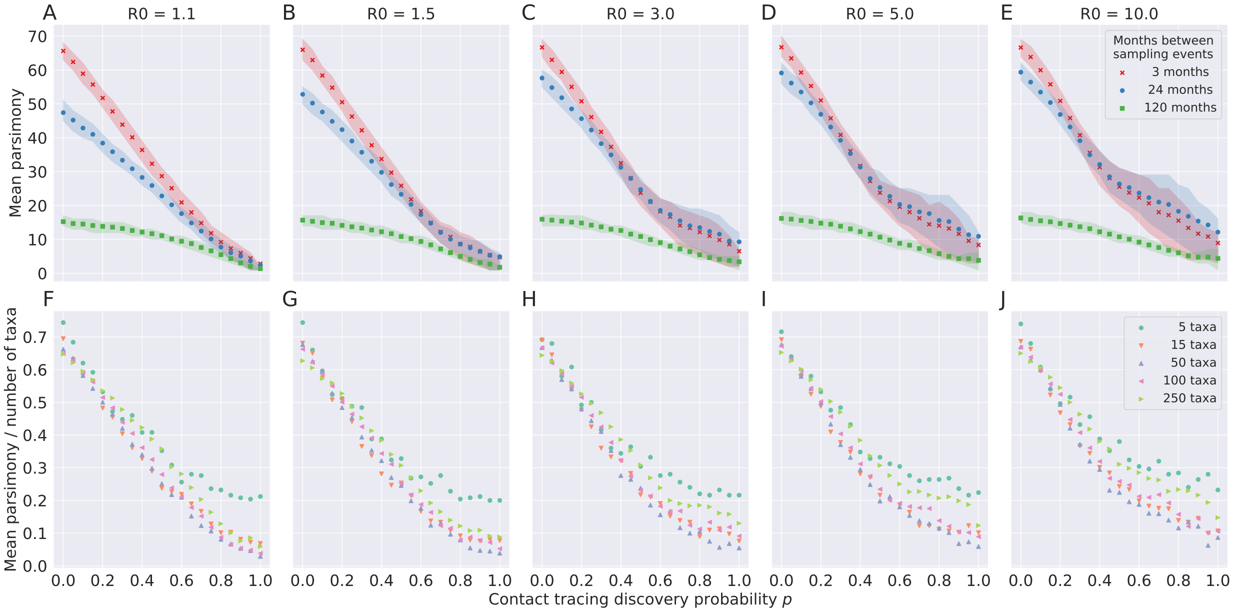


Fig. S4: Parsimony distributions are strongly related to contact tracing performance $p$ and weakly with sample size. The strength of the correlation is primarily dependent on $R_0$ when contact tracing is good. In A-E, a sample of 100 taxa (pathogen sequences from 100 infected hosts) is obtained at the end of the exponential growth phase, and compared against another sample of 100 taken 3, 24, or 120 months later. The shaded region denotes the symmetric inner 50% of the data. In F-J, the experiment is repeated for the 3-month interval between samples, however the sample size is varied. The parsimony score is normalized against the number of taxa in the sampled tree (two times the reported sample size above). Sampled individuals are removed from the population.
